# *Pseudomonas* species from Antarctica and other environments host diverse genes encoding structurally conserved polyhydroxyalkanoate synthases linked with different mobile genetic elements

**DOI:** 10.1101/2025.03.12.642780

**Authors:** Amelia Cox-Fermandois, Camilo Berríos-Pastén, Carlos Serrano, Patricio Arros, Ignacio Poblete-Castro, Andrés E. Marcoleta

## Abstract

Polyhydroxyalkanoates (PHAs) are industrial microbial biopolymers offering a sustainable and biodegradable alternative to petroleum-based plastics. The PHA polymerization process is primarily mediated by PHA synthases (PhaCs), which determine the molecular and physical properties of the synthesized polymers. Previous reports have described Antarctic *Pseudomonas* isolates with unique PhaCs and capabilities for PHA production. However, the genes encoding PhaCs in *Pseudomonas* from Antarctica and other environments have not been investigated systematically. Here, we studied the diversity and phylogenetic distribution of phaC genes in 186 *Pseudomonas* species, including 33 isolates from Antarctica. Most species encode two class II PhaCs, with some displaying additional class II and class I enzymes, especially in Antarctic isolates. Different PhaC subclasses were proposed based on this diversity. Some phaC genes are in putative genomic islands, phages, plasmids, or close to insertion sequences, supporting their acquisition by multiple routes of horizontal gene transfer. Remarkably, the Antarctic strain *P. frigusceleri* MPC6 harbors five PhaCs, including one from a potential novel class. These findings underscore the unique attributes and potential use of Antarctic *Pseudomonas* for biopolymer production. Future research is essential to elucidate the enzymatic properties of this underexplored PhaC diversity.

**Impact Statement:** A few *Pseudomonas* strains have been widely used to produce polyhydroxyalkanoates (PHAs), biopolymers constituting a sustainable alternative to petroleum-based plastics whose synthesis and properties primarily rely on PHA synthase enzymes (PhaCs). We showed that less-studied *Pseudomonas* species, especially from Antarctica, host a remarkable diversity of PhaCs, some likely acquired by horizontal transfer, including a potential novel class with yet-to-be-explored functional features and biotechnological potential.

## INTRODUCTION

Antarctica remains one of the world’s least explored regions, having some of the coldest, driest, and most chemically extreme terrestrial environments for life. Despite these conditions, it hosts diverse prokaryotes that have adapted and thrive, evolving remarkable metabolic features (Holmberg and Jørgensen, 2023; Servettaz et al., 2023). Previous works have shown a strong presence of *Pseudomonas* species in soils of the Antarctic Peninsula and surrounding islands, part of them showing phenotypic traits and producing metabolites with high biotechnological potential, including polyhydroxyalkanoates (PHAs) (Correa and Abreu, 2020; Marcoleta et al., 2022; Orellana-Saez et al., 2019; Ramasamy et al., 2023).

PHAs are microbial hydroxyalkanoic acid molecules linked by ester bonds. These polymers serve as carbon and energy reserves for the cells, present as insoluble granules that provide resistance against freezing, high pressures, and toxic compounds (Możejko-Ciesielska and Kiewisz, 2016; Obruca et al., 2018). PHAs are industrial polyesters displaying properties similar to conventional plastics, such as polypropylene and polystyrene, but are fully biodegradable in soil and water, and can be derived from renewable sources (Rosenboom et al., 2022). The main bacterial genera known for producing PHAs are *Pseudomonas*, *Cupriavidus*, *Bacillus*, *Ralstonia*, and *Alicagenes* (Pawar et al., 2023). In particular, *Pseudomonas* strains have been industrially exploited to produce PHAs, primarily using fatty acids as carbon substrates (Borrero-de Acuña and Poblete-Castro, 2023).

Several *Pseudomonas* species have been described as natural producers of different PHAs (Poblete-Castro et al., 2012; Prieto et al., 2016). In these bacteria, PHA synthesis depends on a well-studied gene cluster composed of two operons, *phaC1ZC2D* and *phaFI*, encoding six key proteins: two PHA polymerases (PhaC1 and PhaC2) responsible for the synthesis of medium-chain-length (*mcl*) PHAs, a depolymerase (PhaZ), two regulatory proteins (PhaD and PhaG), and two phasins (PhaI and PhaF) related to the formation of the PHA granule inside the producing cells (Mozejko-Ciesielska et al., 2019). PhaCs are critical in monomer polymerization, controlling PHA monomeric composition and molecular weight (Neoh et al., 2022). The PhaC enzymes can be classified into four groups (I to IV) based on their substrate specificity, constituent subunits, and the aminoacidic sequence (Rehm, 2003). In *Pseudomonas*, the two PhaCs encoded in the canonical PHA-producing gene cluster belong to class II and have been studied mainly in industrial strains of *Pseudomonas putida* (Poblete-Castro et al., 2012).

We previously reported the isolation and characterization of the Antarctic strains *Pseudomonas* sp. MPC5 and *Pseudomonas frigusceleri* MPC6 from soil collected at Deception Island (South Shetlands, Antarctica). These strains are psychrotolerant extremophiles notable for their capacity to produce high amounts of PHAs from glycerol over a wide temperature range (4 °C to 30 °C), with varying composition but with similar PHA yield (Orellana-Saez et al., 2019; Pacheco et al., 2019). Moreover, the MPC6 strain produced a PHA composed mainly of short-chain length (*scl*) (C3-C5) monomers and blend polymers comprising both *scl* and *mcl*-monomers, being highly different from the typical PHAs synthesized by *Pseudomonas* bacteria, corresponding primarily to *mcl*-PHAs. Furthermore, MPC6 genome analysis indicated that this strain encodes five PhaCs from different classes, which could account for the distinctive PHA produced by this strain. In this line, there is little information regarding the PHA synthesis properties of other Antarctic *Pseudomonas* species and the genetic basis for this trait. In fact, there is a lack of systematic studies on the presence and phylogenetic distribution of PhaC proteins in different *Pseudomonas* species from Antarctica and other geographical locations.

The market positioning of PHA has been restricted by the high production costs. Strategies to reduce production expenses and enhance desired properties include the identification of new microbial strains that exhibit robust PHA accumulation with novel monomeric composition and suitable molecular weights for thermoforming (Neoh et al., 2022). Evaluating the diversity and distribution of PhaCs in different bacteria, particularly those prolific in PHA production, such as *Pseudomonas* spp., is an important step in that direction. This study examined the diversity, classification, phylogenetic distribution, structural features, and possible horizontal transference of PhaC genes in Antarctic *Pseudomonas* compared to strains of this genus isolated across the globe. Our results provide a more representative and insightful panorama of the PhaC content in species of this genus, identifying an array of uncharacterized PhaCs, and highlighting the PHA synthesis potential among *Pseudomonas* from Antarctica.

## MATERIALS AND METHODS

### *Pseudomonas* species genome database construction

A genome dataset from different species of the *Pseudomonas* genus was constructed, including reference and complete genomes from the NCBI database classified as “*Pseudomonas*” (NCBI Taxonomy ID 286) and draft and complete genomes from previously published Antarctic *Pseudomonas* strains. This initial dataset was evaluated regarding assembly quality using CheckM (Parks et al., 2015) and taxonomic classification using the Genome Taxonomy Database (release 214) and its associated software GTDB-Tk (Chaumeil et al., 2022). Genomes showing completeness ≤90%, contamination ≥10%, or that could not be classified or did not belong to the genus *Pseudomonas* were filtered out. Eight hundred fifty-nine genomes met the mentioned inclusion criteria and were considered for downstream analyses (Table S1).

### *Pseudomonas* spp. phylogenomic analysis

The whole genome phylogenetic trees (complete and Antarctic-only datasets) were constructed based on mash distances using Mashtree v1.2.0 (Katz et al., 2019) with the default parameters and 1000 bootstrap iterations. The trees were formatted using the iTOL platform v6 (Letunic and Bork, 2021), incorporating the *phaC* gene presence maps and the GTDB species assignment. Pairwise average nucleotide identity (ANI) calculations were performed with fastANI (Jain et al., 2018).

### Genomic search for *phaC* genes, classification, and conserved domain analysis

Eleven PhaC reference proteins covering the four known PhaC classes were downloaded from the NCBI database (Table S2) and used to search for *phaC* genes in the *Pseudomonas* genome dataset. In addition, we included as a reference the PhaC3 protein identified previously in the Antarctic *Pseudomonas frigusceleri* MPC6 strain (QCY11757.1), which was suggested to belong to a novel PhaC class (Orellana-Saez et al., 2019). The search for PhaCs among the unannotated genomes was done with BLASTX (Gish & States, 1993) with the default parameters. The output tables with the sum of hits for all the genomes were analyzed to examine the percentage identity and coverage frequency distribution. The *phaC* genes were assigned to a PhaC class based on the best hit. Identity and coverage thresholds for positive hits were defined based on the distribution observed for each PhaC class. For class-II PhaCs, 60% identity and 80% coverage were used, while for class I, we kept the 80% coverage threshold but lowered the identity to 50% as more sequence diversity was noticed. The amino acid sequences translated from the *phaC* genes were extracted from the genomes using custom Python scripts. Conserved domains were identified by comparison with the Conserved Domain Database v2023 (Wang et al., 2023) using BLASTp (Altschul et al., 1990).

### Multiple sequence alignment and construction of phylogenetic trees of the extracted PhaCs

Multiple sequence alignments were performed using MAFFT v7 (Katoh and Standley, 2013). Before phylogenetic inference, non-informative blocks and gaps were removed from the alignment using BMGE v1.12_1 (Criscuolo and Gribaldo, 2010). The cleaned alignments were used as input for maximum likelihood phylogenetic tree inference with IQ-tree v2.2.5 (Minh et al., 2020). IQtree runs were conducted with 10,000 ultrafast bootstraps (UFBoot) (Hoang et al., 2018) and the-bnni option to reduce the risk of overestimating UFBoot supports when the data severely violates the model. ModelFinderPlus (Kalyaanamoorthy et al., 2017) was used to accurately select the best amino acid substitution model for each dataset: Q.pfam+R3 for class-I PhaCs and Q.plant+I+R7 for class-II PhaCs. 139,808 was used as a seed for random number generation. Then, the iTOL platform v6 (Letunic and Bork, 2021) was used to mid-point root and format the obtained trees, incorporating relevant metadata as the taxonomic species and PhaC subclass.

### PhaC 3D structure analysis

The 3D protein structure of selected PhaC proteins was predicted using ESMfold_v1 (Lin et al., 2023), executed with the default number of recycles (4). Structural alignments based on the predicted 3D structures were performed using US-align (Zhang et al., 2022). Protein structure visualization, coloring, and rendering were performed using Pymol (Schrödinger, LLC, 2015).

### Analysis of the phaC genomic contexts

The genomic context of the *phaC* genes in different *Pseudomonas* species was investigated by annotating the genome of selected representatives using PROKKA (Seemann, 2014). Then, 40-kbp upstream and downstream regions were extracted and examined using the Snapgene software (www.snapgene.com) and customized Python scripts. The *phaC* genomic contexts were compared and plotted using Clinker (Gilchrist and Chooi, 2021). Insertion sequences were predicted using ISfinder (Siguier et al., 2006). Direct repeats and integrase-coding genes were identified using BLASTn and BLASTx, respectively.

## RESULTS

### Species and phylogenomic relationships among a diverse *Pseudomonas* genome set to study *phaC* gene content

We investigated the phylogenetic relationships and *phaC* gene content in the genomes of 859 strains encompassing 186 species of *Pseudomonas*, according to the taxonomic classification based on the Genome Taxonomy Database. Thirty-three strains were isolated from Antarctica, while the rest came from different geographic locations and biomes. The most represented species in our dataset were *P. aeruginosa* (331), followed by *P. chlororaphis* (53), *P. fluorescens* (23), *P. avellanae* (20), and *P. putida* (20) (Table S1). Phylogenetic analysis based on whole-genome distance calculations and tree inference using Mashtree revealed a highly diverse population with deep branching lineages, except for *P. aeruginosa,* which showed a reduced genetic variability (Figure 1A). In this sense, *P. aeruginosa* showed an ANI median=98.95% and standard deviation=0.94, while it was slightly lower in *P. avellanae* (97.42±2.01%) and *P. chlororaphis* (95.89±2.57%), the latter being in the edge of currently accepted ANI species boundaries (Jain et al., 2018). Conversely, the ANI values were notably lower among *P. fluorescens* (83.58±3.68) and *P. putida* (85.94±4.33%), underlining the high genomic diversity in these lineages, likely corresponding to species complexes. In agreement, *P. fluorescens* and *P. putida* genomes were spread in different clades in our phylogenetic tree. This evidence agrees with *P. aeruginosa* and *P. avellanae* being more adapted and restricted to animal and plant host-associated niches, while *P. fluorescens* and *P. putida* are found in a broader range of environments. This high genomic variability anticipates a potentially wide and unexplored variety of PhaCs and PHA synthesis capabilities across different *Pseudomonas* species.

**Figure 1.**
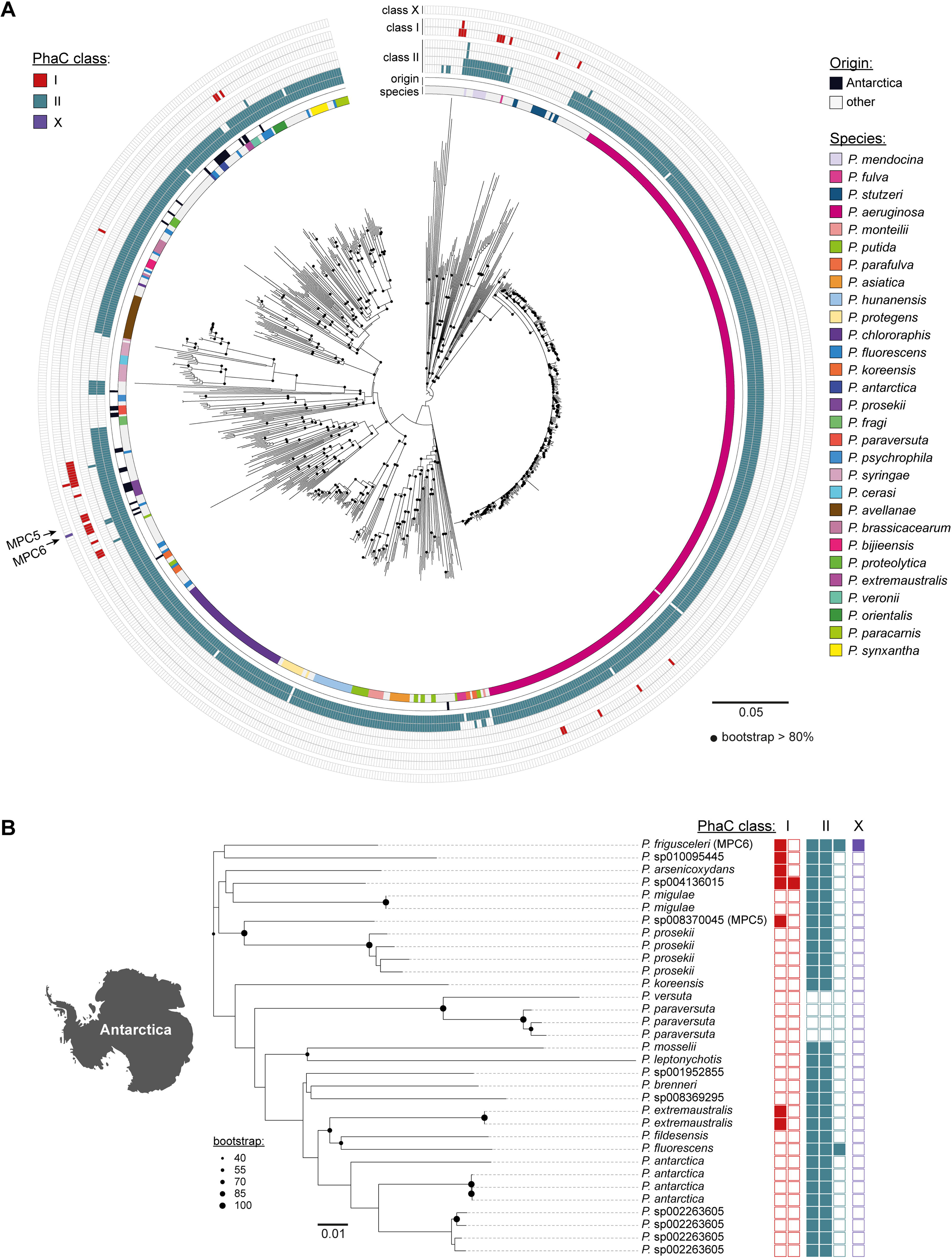
Phylogenomic relationships and PhaC gene content across different Pseudomonas species revealed core and sporadic PhaC enzymes, especially in Antarctic strains. A: Phylogenetic tree constructed based on whole-genome distance calculations using mashtree, based on 859 genomes encompassing 186 *Pseudomonas* species. 1000 bootstrap iterations were performed. The tracks show the species, Antarctic origin (or not), and the presence of one or more class-I, class-II, and class-X PhaCs. B: Mashtree showing only the *Pseudomonas* strains isolated from Antarctica.

### Diversity and distribution of genes encoding PhaCs among *Pseudomonas* species

Using BLASTX and a set of reference PhaCs covering the four known classes (I-IV) (Table S2), we searched our *Pseudomonas* genome set for genes encoding PhaCs. Class-III and class-IV PhaCs were ruled out, as only non-specific hits with less than 38% identity and low coverage were obtained. Conversely, varying genes encoding class-I and class-II PhaCs were observed.

Despite the species diversity, most strains (738) had genes encoding two class-II PhaCs (Figure 1A) located in close genomic proximity and thus likely correspond to these encoded in the canonical cluster found in *Pseudomonas*, commonly known as PhaC1 and PhaC2. For simplicity and consistency when naming PhaCs from other classes, we will refer to these well-known PhaCs from classes II.1 and II.2, respectively.

Remarkably, six of these strains had a third and one a fourth class-II PhaCs (*P. mandelii*; *P. laurylsulfatiphila*; *P. silesiensis*; *P. fluorescens*; *P. frigusceleri*, and *P. lalkuanensis*). These additional *phaC* genes were in genomic regions distinct from the two genes found widely present across the genus and were sporadically detected in some isolates from various species. For instance, from the three *P. mandelii* genomes, only one encoded a third class-II PhaC. Similarly, from the 23 *P. fluorescens* genomes, only one carried a third PhaC. This sporadic presence across different species suggests these enzymes are linked to mobile genetic elements facilitating their dissemination through horizontal gene transfer.

Some genomes had PhaC from other classes, which was reported as uncommon in *Pseudomonas* (Neoh et al., 2022). In this concern, 38 genomes encoded one class-I PhaC, including *P. aeruginosa* and 23 other species, while two had a second class-I PhaC (*P.* sp004136015 and *P. oryzae*). Overall, the class-I PhaCs showed a sporadic presence in some species, including *P. aeruginosa* (5/331 strains), *P. stutzeri* (2/12), *P. extremaustralis* (2/3), *P. veronii* (1/4) and *P. fluorescens* (2/23). Contrastingly, we found some strains that completely lacked genes encoding PhaCs. In general, this absence correlated with the species. For example, all the *P. syringae* and *P. stutzeri* strains in this study lack PhaC-coding genes. Unfortunately, most sequenced *Pseudomonas* genomes in databases correspond to *P. aeruginosa*, leaving other species much less represented. This caveat prevents a more exhaustive analysis of the PhaCs presence within these species.

A second phylogenetic tree was constructed using genomes from Antarctic *Pseudomonas* only to investigate whether they show distinctive patterns in the phaC content (Figure 1B). Notably, out of the 33 Antarctic *Pseudomonas* isolates encompassing 20 different species, eight encoded extra class-II or class-I PhaCs (Figure 1B). The high diversity of species captured in this relatively small collection of Antarctic *Pseudomonas* genomes is remarkable, suggesting the presence of species yet to be discovered at the genomic level in this continent. Among these, *P. frigusceleri* MPC6 is distinguished for encoding one class-I and three class-II PhaCs, and it is the only strain in our genome collection that encodes a highly divergent PhaC, previously suggested to belong to a novel class (Orellana-Saez et al., 2019), provisionally named class X. Additionally, *Pseudomonas* sp. 004136015 was found to encode two class-I PhaCs, while *Pseudomonas* sp. MPC5 encoded one class-I PhaC besides the typical two class-II enzymes. These findings emphasize the vast diversity of *Pseudomonas* species in Antarctica and their potentially novel attributes related to PHA synthesis. However, the absence of PhaCs in Antarctic strains of *P. versuta and P. paraversuta* indicates that, while PHA production may be beneficial for adaptation to the extreme Antarctic environment, it is not strictly required for the survival of *Pseudomonas* in this adverse ecological niche. How the canonical cluster, present in most other *Pseudomonas* species, was lost in certain lineages remains to be investigated.

### Properties of class-II PhaCs found in *Pseudomonas*

Next, we aimed to characterize the PhaCs identified among our *Pseudomonas* genome set, assessing their diversity within this genus, phylogenetic relationships, and structural features. For this purpose, we inferred a phylogenetic tree based on the multiple sequence alignment of the PhaC sequences, revealing a significant diversity inside class-II enzymes in *Pseudomonas* (Figure 2A). Two principal, well-differentiated clades were observed aligning with the two class-II polymerases typically encoded in the canonical cluster (II.1 and II.2). Several subclades occurred within these groups, which predominantly included sequences from the same or related species, suggesting divergence driven by speciation.

**Figure 2.**
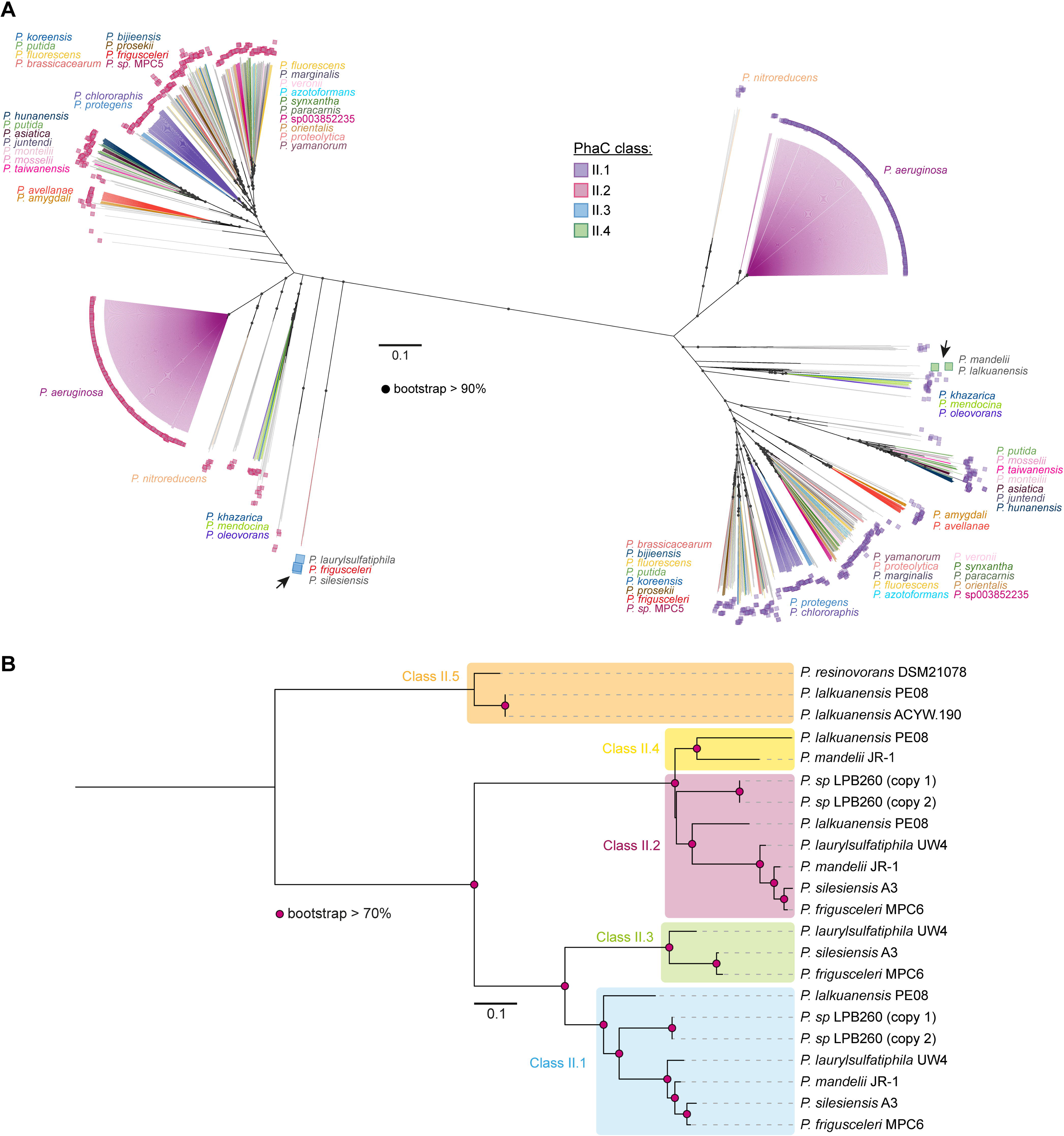
Class-II PhaC enzymes found in different *Pseudomonas* species. A: Maximum-likelihood tree inferred from the multiple amino acid sequence alignment of the PhaC proteins found in the genome collection representing 186 Pseudomonas species. 10,000 ultrafast bootstraps were calculated. The arrows point to sporadic class II.3 and II.4 PhaCs. B: Simplified tree showing representatives of the four PhaC sub-classes found in the genome set described here (II.1-II.4), plus a fifth highly divergent class-II PhaC found in recently published genomes from *P. resinovorans* and *P. lalkuanensis* (class II.5).

The additional class-II PhaCs present in *P. fluorescens* UKR4 and *P. sp.* LPB260 genomes clustered within clades II.1 and II.2 and corresponded to 100% identical copies of the respective genes from the canonical cluster. In this concern, the UKR4 assembly is highly fragmented (889 contigs) and has relatively high contamination (8.88% as evaluated by CheckM software), while the LPB260 genome showed a 3 Mbp duplicated region. Thus, these extra class-II PhaC gene copies likely arose from assembly errors and thus do not represent actual cases of additional class-II PhaCs.

We observed a more divergent clade of sporadic class II PhaCs related to class II.1, present in *P. laurylsulfatiphila* UW4, *P. frigusceleri* MPC6, and *P. silesiensis* A3. Despite being present in different species, the A3 and MPC6 PhaC shared 98% amino acid identity, while UW4 showed ∼86% identity with the formers. These enzymes had 67-69% identity with the canonical class II.1 PhaC from the same strain and thus correspond to different class-II enzymes (named class II.3).

Two strains encoded sporadic PhaCs more closely related to class II.2 enzymes, namely, *P. mandelii* JR-1 and *P. lalkuanensis* PE08. These PhaCs shared 77% identity, and showed 77% and 74% identity with the II.2 PhaC from the same strain, respectively. Thus, these genes encode proteins that are different from class II.2 (named II.4).

When comparing the class II PhaCs found in our genome dataset with those in the NCBI database, we noticed the presence of further highly divergent class II enzymes in three strains, *P. lalkuanensis* PE08, *P. lalkuanensis* ACYW190, and *P. resinovorans* DSM21078. Strains ACYW.190 and PE08 shared 100% identity while showing 88.94% identity with DSM21078. Diverging class-II PhaC shared 45-48% identity with other class II and 33-35% with Class-I PhaCs. Thus, they were assigned to subclass II.5. A simplified phylogenetic tree including representatives from the different class II subclasses is depicted in Figure 2B.

Sequence identity comparison among class II.1-II.5 PhaC representatives showed congruent grouping with the defined subclasses, as well as the invariable presence of PhaC (COG3243), PhaC_N (pfam07167), and PHA_synth_II (TIGR01839) domains (Figure 3A).

**Figure 3.**
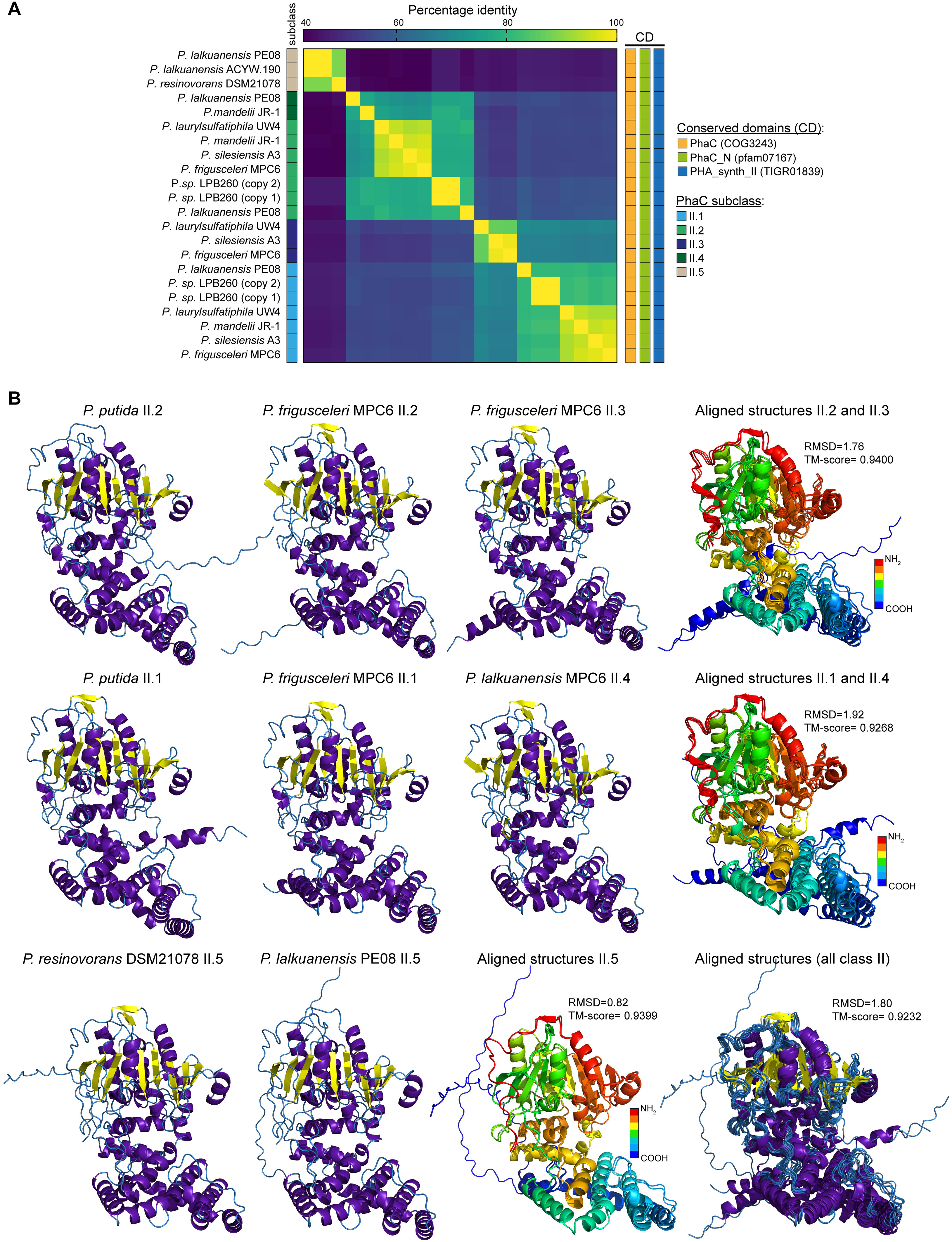
Amino acid sequence identity and structural features of the class-II PhaC enzymes found in different *Pseudomonas* species. A: Heatmap showing the pairwise identity between enzymes from the five sub-classes found. The presence of conserved domains is indicated at the right of the heatmap. B: predicted 3D structure of representatives of the different class-II PhaC sub-classes. Structural alignments revealed high conservation between all the structures, as revealed by the template modelling (TM) score and the root mean square deviation (RMSD) values.

PhaC structure prediction and comparison showed notable conservation of the global protein topology and the main secondary structure elements, including the N-terminal alpha-helices composing the oligomerization surface and the beta-strands forming the substrate-binding domain (Figure 3B). Class II.2 PhaCs of *P. frigusceleri* MPC6 show high structural similarity with the corresponding reference proteins of *P. putida*, as indicated by their TM-score: 0.91344 and 0.94042, respectively, which correlated with the relatively high amino acid sequence identities between these protein pairs (81.57% and 71.96%). Moreover, II.2 and II.3 enzymes also share high structural similarity (Tm=0.94; RMSD=1.76), as well as class II.2 and II.4 (TM-score=0.9268; RMSD=1.92). Furthermore, even the highly divergent class II.5 PhaC, exhibiting nearly 40% sequence identity with classes II.1-II.4, showed high structural conservation. This high structural conservation across class II is further supported by a TM-score of 0.9232 and an RMSD of 1.8 upon aligning the representatives from all the subclasses. Nevertheless, most variations occur in the N-terminal half, resulting in a different number and disposition of alpha helixes connected by flexible loops. Additional studies are needed to elucidate the impact of these subtle structural variations and the high amino acid sequence diversity on these PhaC’s functional features.

### Features and phylogenetic relationships of *Pseudomonas* spp. class-I and class-X PhaCs

Phylogenetic inference for class-I PhaCs revealed several well-defined clades, from which we identified eight class-I PhaCs subclasses (Figure 4A). Clade I.1 included proteins of 566-567 amino acids sharing 88-99% identity, found in twelve different species. Clade 2 included proteins of 567 amino acids sharing 87-94% identity, present in four different species. Clade 3 included proteins of 566 amino acids sharing 89-100% identity, all of them from *P. aeruginosa*. Clade 4 included proteins of 566-568 amino acids sharing 99-100% identity, found in seven species.

**Figure 4.**
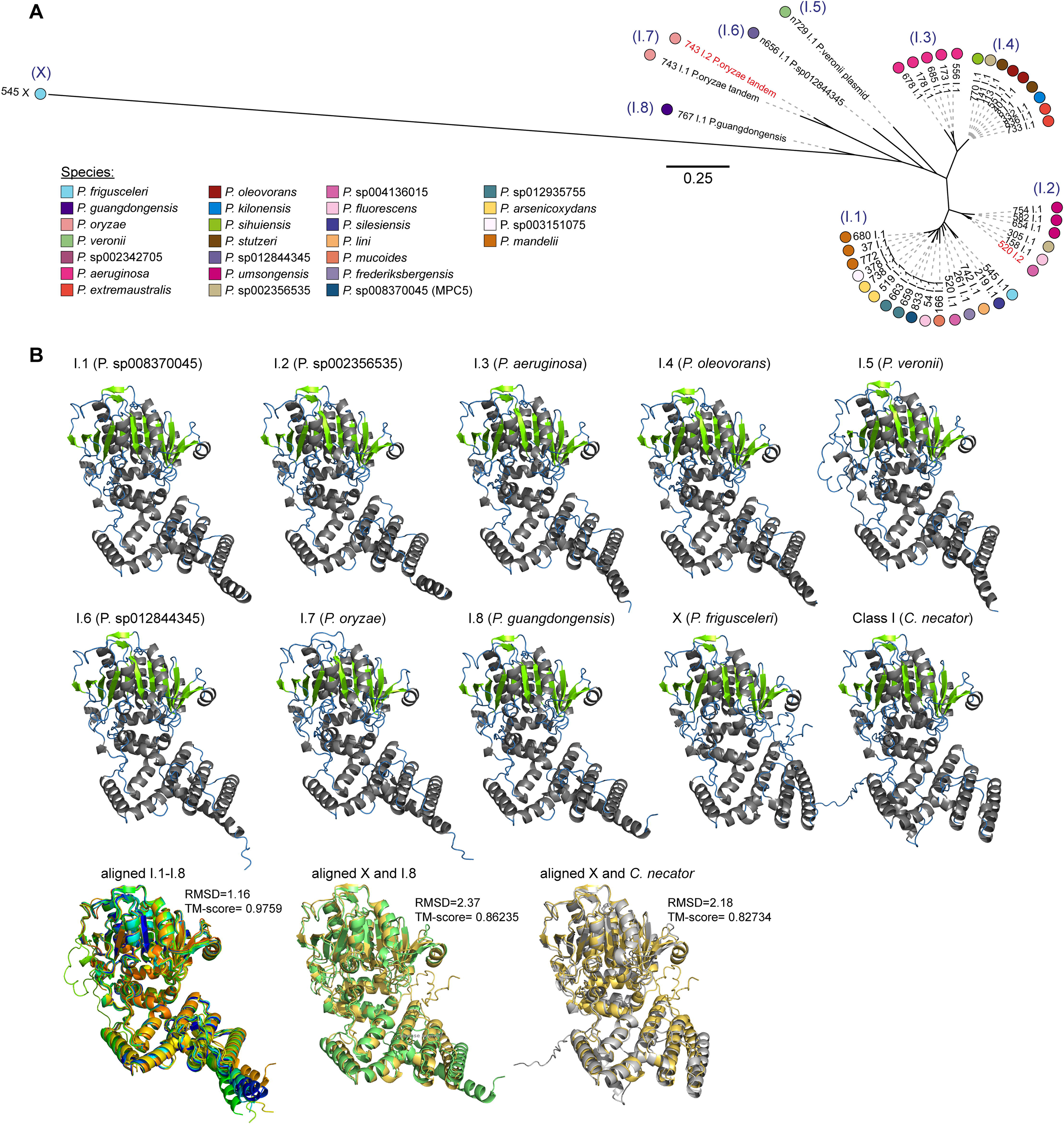
Diversity and structural features of Class-I and class-X PhaC enzymes found in different *Pseudomonas* species. A: Phylogenetic tree inferred from the multiple amino acid sequence alignment of 42 class-I and class-X PhaC proteins found in 25 *Pseudomonas* species. Sub-classes I.1-I.8 were defined based on the clustering. Bootstrap values were omitted for more clarity. B: predicted 3D structure of representatives of the different class-I PhaC sub-classes. In the single (non-superimposed) structures, beta-sheets are indicated in green, while alpha helices are in gray. Structural alignments revealed high conservation between all the structures, as revealed by the TM-score and the RMSD values. Class-I PhaC from *Cupriavidus necator* is shown as a reference. Superposed aligned structures show each protein in a different color.

More divergent class-I PhaCs included the one found in *P. veronii* and P. sp012844345, which shared 66-71% and 67-69% identity, respectively, with the enzymes from the clades 1 to 4. Meanwhile, the two class I PhaCs found in *P. oryzae* shared 83% identity, showing 59-70% identity with the rest of the class-I PhaCs. The PhaC from *P. guangdongensis* showed 63-67% identity with the other Class-I PhaCs. Contrastingly, the class-X PhaC, although possessing a class-I PhaC conserved domain, was highly divergent, sharing 30-36% identity with the class-I enzymes included in the analysis. This high divergence from other class-I PhaCs strongly suggests that this enzyme belongs to a different class.

The analysis of conserved domains revealed that all the class-I and the class-X PHA synthases shared the PhaC domain (COG3243), the PHA_synth_I domain (TIGR01838), and the N-terminal PhaC domain (pfam07167) with the class-I reference protein from *Cupriavidus necator* (Figure S1).

We aimed to explore the structural properties of the class-I PhaCs found in *Pseudomonas* species. Thus, we predicted and compared the 3D structure of class-I and class-X PhaC representatives, including the *C. necator* class-I PhaC crystal structure as a reference (Figure 4B). Remarkably, despite the limited amino acid identity between some of them, class I.1-I.8 PhaCs exhibited highly similar secondary and tertiary structures, with a TM-score of ∼0.98 and an RMSD of 1.16. Moreover, class-X PhaC also demonstrated a highly similar 3D structure (TM-score: 0.86235; RMSD: 2.37) with class-I.8 PhaC, which is the closest in the phylogenetic tree. Notably, this potentially novel-class PhaC showed an ordination of the N-terminal alpha-helices more similar to the *C. necator* PhaC than the class I PhaCs found in *Pseudomonas*.

Given the remarkable divergence of class-X PhaC with the other known PhaCs, we aimed to investigate if other published genomes from *Pseudomonas* not included in our database, or maybe other bacterial lineages, encode one or more class-X PhaCs. BLAST analysis against the non-redundant protein database revealed that 99-100% identical proteins were found encoded in the genome of *P. silesiensis* Phyllo_186 (MEX5686240.1), *Pseudomonas* sp. VI4.1 (WP_128871497.1), and *P. fluorescens* PS928 (WP_150786187.1). Although limited in number, this evidence supports the existence of this protein in other *Pseudomonas* strains. Hits in species from other genera showed 69% or less identity, suggesting that this protein is mainly restricted to *Pseudomonas*.

### Genomic context of class II PhaC-coding genes in *Pseudomonas*

We examined the genomic context of the PhaC coding genes in representatives of different *Pseudomonas* species. As expected, II.1 and II.2 PhaCs were found along with the other genes composing the canonical PHA production cluster: *phaZ* (PHA depolymerase), *phaD* (transcriptional regulator), and the *phaF* and *phaI* genes encoding granule-associated phasin proteins (Figure 5). Notably, *P. paraversuta* and *P. syringae* showed a deletion encompassing the *phaC1*, *phaZ*, and *phaC2* genes. The cause or mechanisms mediating this deletion could not be established since no signs of mobile genetic elements were found in the surroundings of these genes.

**Figure 5.**
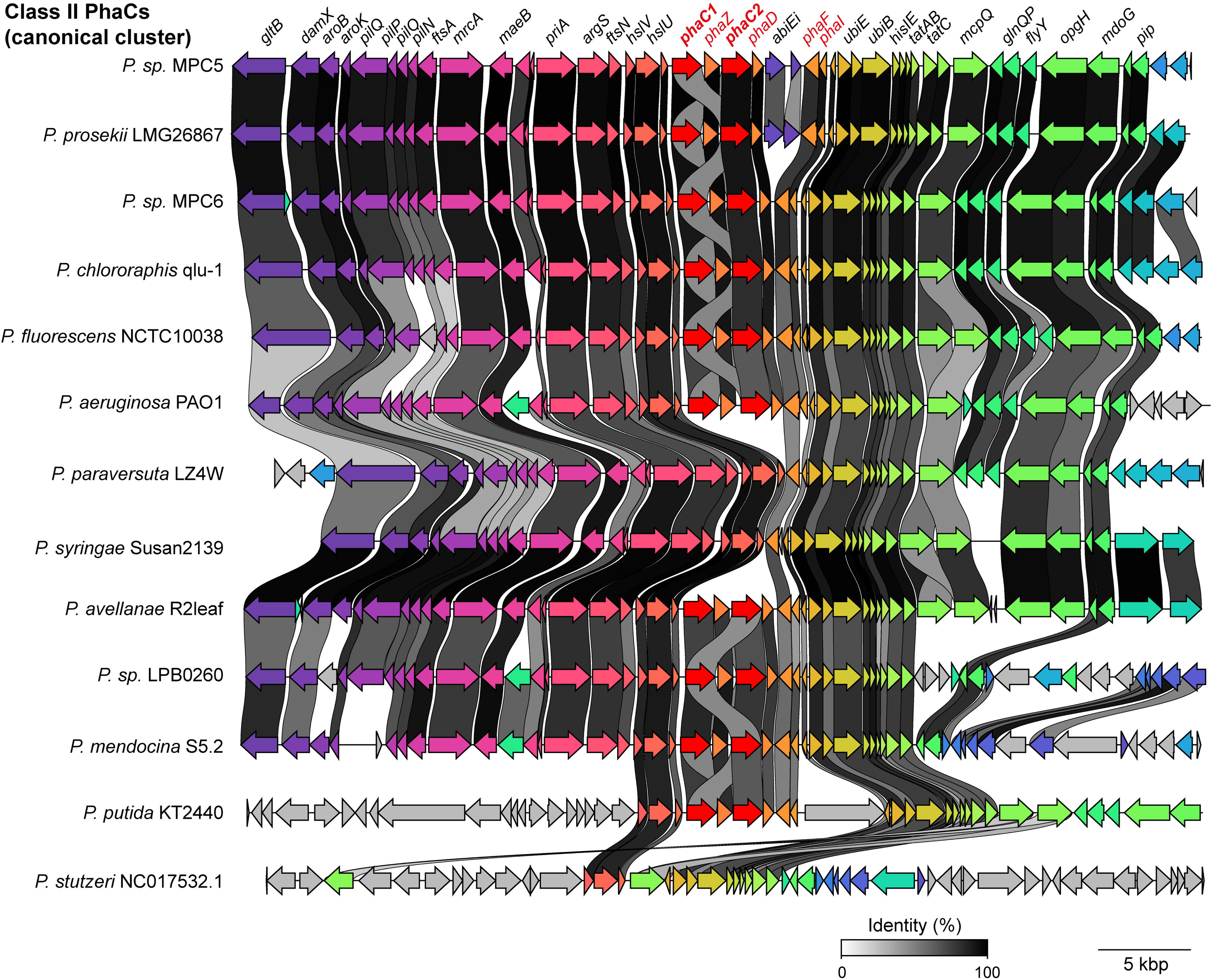
The canonical class-II PhaC gene cluster and its genomic context are highly conserved in different *Pseudomonas* species. Sequence alignment and visualization using the tool Clinker of the genes encoding the two class-II PhaCs found in most *Pseudomonas* species (PhaC1 and PhaC2) and their genetic environment. Notably, PhaC genes are absent in *P. paraversuta, P. syringae and P. stutzeri*.

Regarding the sporadic genes encoding class II PhaCs, we found them in variable genomic contexts depending on the PhaC subclass. Class II.3 *phaC* in *P. laurylsulfatiphila* UW4 was found inside a predicted genomic island (GI) integrated into a Ser tRNA gene and flanked by 72-bp direct repeats that include the 3’ end of the tRNA gene (Figure 6A), as typically found in GIs from *Pseudomonas* and other Gram-negative bacteria (Juhas et al., 2009), although lacked predicted integrase-coding genes. Additionally, this GI included the *phaZ* gene (depolymerase) from the canonical cluster and genes with functions potentially linked to PHA production and lipid metabolism, including a putative acyl-CoA dehydrogenase, a lipid-transfer protein, a 3-oxoacyl-ACP reductase (*fabG*), an arylsulfatase (*atsA*), a choline oxidoreductase, and a long-chain fatty acid-CoA ligase (Figure 6A). Moreover, supporting the finding of this genomic island, we found high variability in the equivalent Ser tRNA locus in other *P. laurylsulfatiphila* strains lacking II.3 *phaC* genes, including a prophage in *P. laurylsulfatiphila* B21-025 or the absence of mobile elements in *P. laurylsulfatiphila* AP316 (Figure 6A). Conversely, the lack of additional strains from *P. frigusceleri* and *P. silesiensis* species prevented further inferences regarding the mobility of the chromosomal region encoding the *phaC* gene. In MPC6, an IS53-like transposase was found close upstream of *phaC* (Figure 6A), suggesting the role of insertion sequences in mobilizing this gene. Noteworthy, genes surrounding II.3 *phaC* in MPC6 included several linked to arsenic metabolism, transporters, and ATPases.

**Figure 6.**
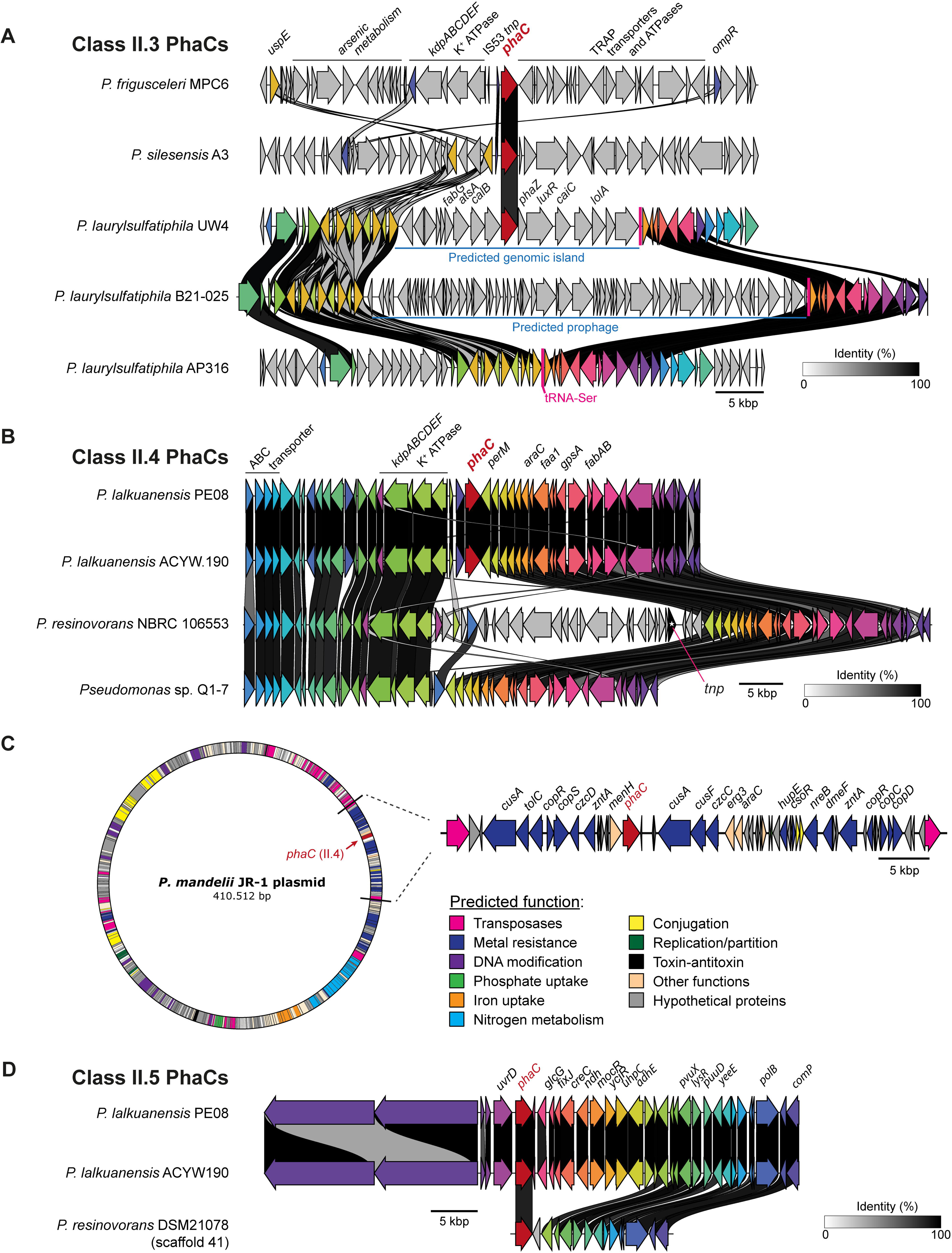
Genomic context of sporadic class-II PhaC genes in different *Pseudomonas* species, some showing association with mobile genetic elements. A: Class II.3 PhaCs in *P. frigusceleri*, *P. silesiensis*, and *P. laurylsulphatiphila*. In the latter species, the phaC gene was found inside a putative genomic island. *P. laurylsulphatiphila* strains B21-025 and AP3 16 lacking II.3 PhaC genes were included for comparison. B: Class II.4 PhaCs found in *P. lalkuanensis*. Strains *P. resinovorans* NBRC 106553 and *P. sp.* Q1-7 lacking class II.4 PhaC genes were included for comparison. C: Plasmid from *P. mandelii* carrying a class-II PhaC gene and genes for metal resistance, phosphate and iron uptake, and nitrogen metabolism. D: genes encoding PhaCs from the highly divergent class II.5. Arrows in black with a white star inside represent genes encoding transposases.

Class II.4 PhaC genes in *P. lalkuanensis* PE08 and ACYW.190 were found in a variable genome region compared with strains from related species (Figure 6B). No tRNA-coding genes, direct repeats, or integrase genes were found in this case. In an equivalent chromosome position, *P. resinovorans* NBRC106553 harbored a 25-kbp insertion encoding a transposase (among other genes) and lacking *phaC*, while in *P. sp*. Q1-7 (lacking class II.4 PhaC genes, included for comparison), we found no integrations in this region. Of note, the region immediately upstream of the *phaC* gene in *P. lalkuanensis,* including the *kdpABCDEF* genes and a K^+^ ATPase, was similar to the class II.3 *phaC* upstream region found in *P. frigusceleri* MPC6. Meanwhile, in *P. mandelii* JR-1, the class-II.4 PhaC gene was found in a 410-kbp potentially conjugative plasmid (Figure 6C) encoding several genes for metal resistance, phosphate uptake, and nitrogen metabolism, among others.

The highly divergent class II.5 PhaC found in *P. lalkuanensis* and *P. resinovorans* could not be associated with any mobile element, mainly due to the lack of genome-sequenced strains to compare (Figure 6D). This *phaC* gene is not accompanied by other genes from the canonical cluster or genes with potential functional relatedness.

### Genomic context of class I PhaC-coding genes in *Pseudomonas*

Class-I PhaC genes were found in highly variable genomic contexts and thus are likely associated with mobile genetic elements. In different species, including *P. mandelii*, *P. mucoides*, *P. fluorescens*, *P. silesiensis*, and *P. frigusceleri*, among others, class I.1 PhaC genes were found in putative genomic islands integrated into selenocysteine (Sec) tRNA genes (Figure 7A). This integration site was in an equivalent chromosome position across these species, showing a highly conserved upstream genomic context and variable downstream genes, supporting it as a hotspot for the traffic of mobile elements targeting. In these putative GIs, *phaC* was almost invariably accompanied by genes *phaR* (transcriptional regulator), *phaP* (phasin), DUF3141, *phbB* (3-oxoacyl-[acyl-carrier-protein] reductase), *paaJ* (acetyl-CoA C-acetyltransferase), *ycaC* (nicotinamidase-related amidase), and *oprD* (porin). Also, *perM* (permease), *fixJ* (Two-component response regulator), *menE* (fatty-acyl-CoA synthase), *actP* (cation/acetate symporter), and *caiA* (Acyl-CoA dehydrogenase). Meanwhile, in P. *arsenicoxydans*, the class I.1 PhaC gene was found in a different genomic context, not in the proximity of tRNA genes, although harboring all the genes mentioned above in a similar organization.

**Figure 7.**
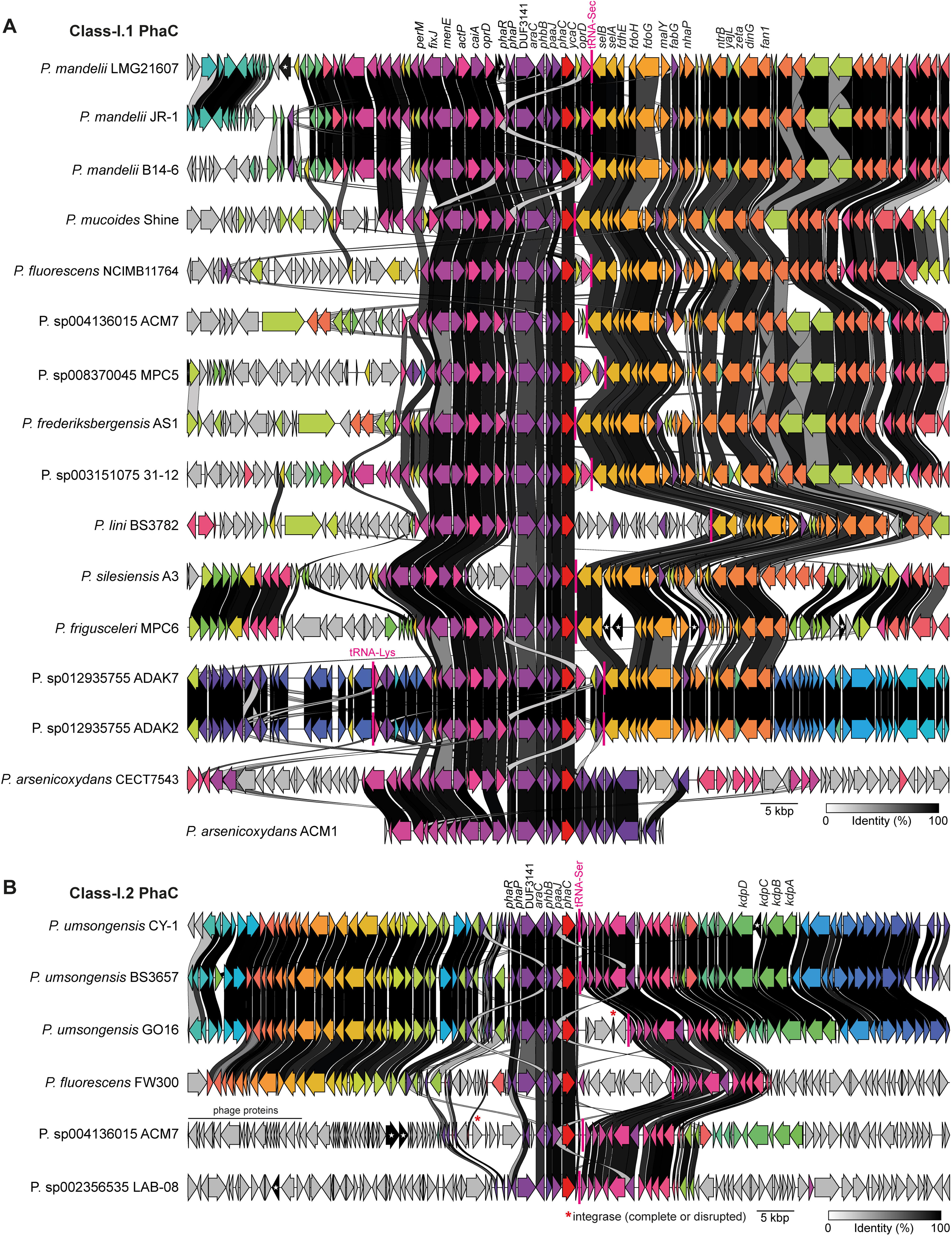
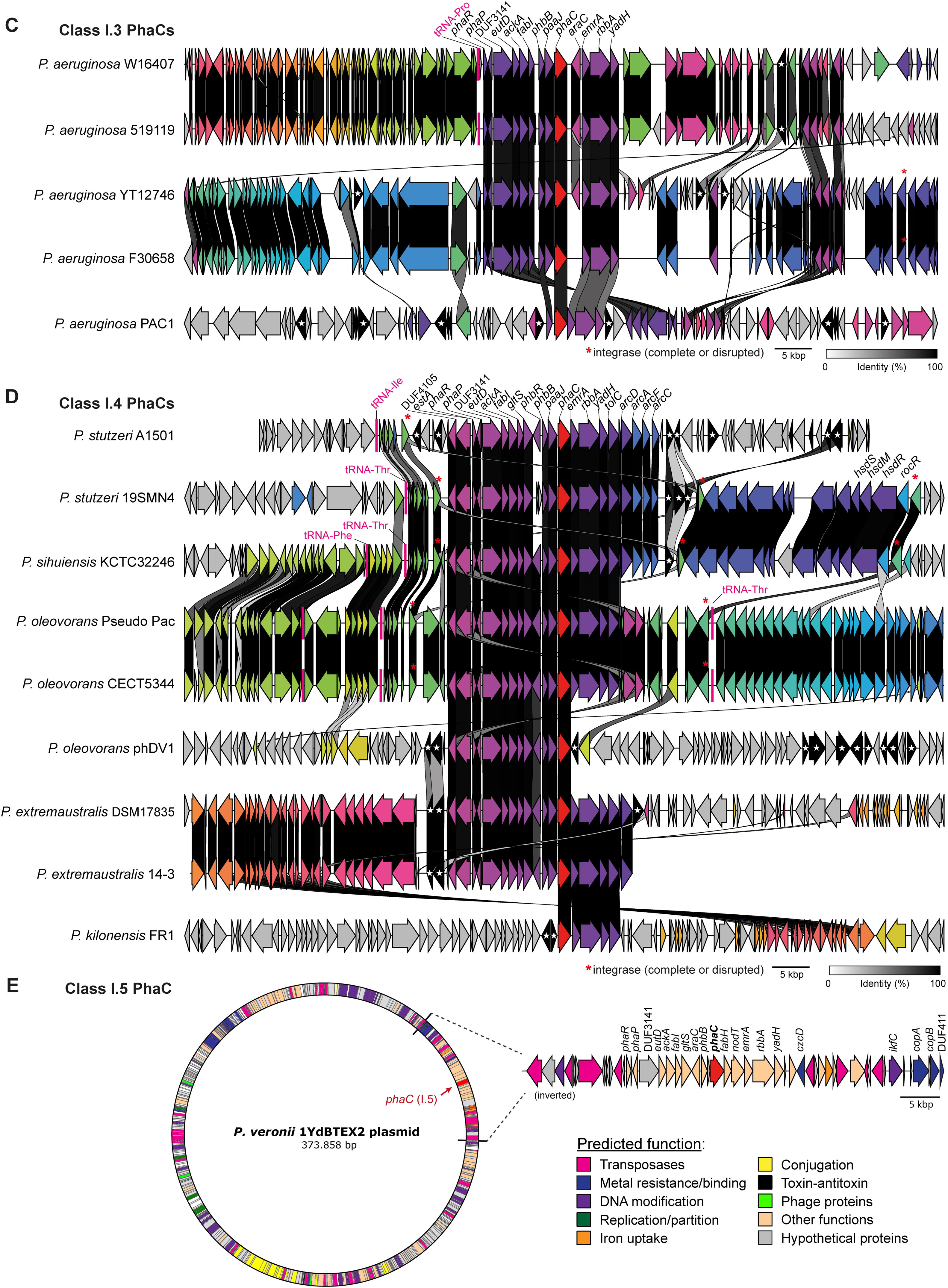
Genes encoding class-I PhaC and its genetic environment in different *Pseudomonas* species. A: Class I.1 PhaCs. B: Class I.2 PhaCs. C: Class I.3 PhaCs. D: Class I.4 PhaCs. E: Class I.5 PhaCs. Arrows in black with a white star inside represent genes encoding transposases. Arrows with a red asterisk indicate integrase-coding genes.

Similarly, class I.2 PhaC genes were found in a variable region surrounded by a highly conserved genomic context, mainly in *P. umsongensis* strains, potentially corresponding to genomic islands integrated into Ser tRNA genes, some carrying complete or disrupted integrase genes (Figure 7B). In this case, PhaC genes were accompanied by a more reduced set of genes than class I.1: *phaR*, *phaP*, DUF3141, *araC*,

*phbB*, and *paaJ*. In *P.* sp004136015 ACM7 and *P.* sp002356535 LAB-08 the *phaC* gene was found in highly divergent genomic regions although sharing the tRNA-Ser and its immediate upstream context (genes depicted in shades of pink). Moreover, in the former strain, the *phaC* gene region downstream of the Ser tRNA included a complete phage-like integrase and several structural phage proteins, indicating that phaC genes would also disseminate through prophages infecting *Pseudomonas*. The scarce number of sequenced genomes from some *Pseudomonas* species prevented a more precise delimitation of the putative mobile elements found.

Class I.3 PhaC genes were found inside a putative genomic island integrated into a Pro tRNA gene in *P. aeruginosa* W16407 and 519119, along with *phaR*, *phaP*, DUF3141, *eutD* (phosphate acetyltransferase), *ackA* (acetate kinase), *fabI* (enoyl-ACP reductase), *phbB*, *paaJ*, and the components of an efflux pump *emrA*, *rbbA*, and *yadH* (Figure 7D). Although the mentioned genes surrounding *phaC* and their synteny are preserved, the genomic context surrounding them is markedly different in *P. aeruginosa* YT12746, F30658, and PAC1, with the absence of tRNA genes in all of them, and an inversion and multiple insertion sequences in the latter strain. In addition, complete integrase-coding genes were found downstream of *phaC* in the strains YT12746 and F30658.

Regarding class I.4, found in *P. stutzeri*, *P. suhuiensis*, *P. oleovorans*, *P. extremaustralis,* and *P. kilonensis*, they would be encoded in genomic islands integrated into Thr and Ile tRNA genes, several carrying integrase genes and insertion sequences (Figure 7D and Figure S2). *P. stutzeri* 19SMN4 putative GI (∼69 kbp) was flanked by 50 bp direct repeats encompassing the 3’ end of the Ile tRNA, as typically occurs for genomic islands targeting tRNA genes.

A class I.5 PhaC gene was found in a potentially conjugative 372-kbp plasmid from *P. veronii* 1YdBTEX2, also carrying genes for metal resistance, iron uptake, and some phage-related proteins (Figure 7E). Besides, classes I.6, I.7, and I.8 were found in highly variable genomic contexts, although no evident mobile genetic elements were detected (Figure S3). Despite this variability, we found this enzyme always accompanied by DUF3141, PhaR, PhaP, although variations among strains were noticed. All this evidence points out that at least part of the sporadic class I and class II PhaCs are associated with mobile genetic elements and, thus, subject to horizontal gene transfer.

## DISCUSSION

PhaC enzymes are essential for PHA synthesis, as they catalyze the polymerization and thus determine the *R-3-*hydroxyacyl-CoA monomer composition of the polymer (Neoh et al., 2022). Hence, studying *phaC* gene diversity and properties is crucial for assessing the PHA synthesis potential in diverse bacterial taxa and identifying novel PhaCs that could benefit industrial applications. These new enzymes can potentially reduce production costs and enhance the physical and mechanical properties of the resultant polymers. In this sense, our study evaluated the presence of PhaC genes across 186 *Pseudomonas* species, exploring their capacity for PHA production. This way, we uncovered a group of previously unrecognized PhaCs, pointing out a vastly underestimated *phaC* diversity across this genus, especially in Antarctic *Pseudomonas* isolates.

Despite the gene cluster for PHA production in *Pseudomonas* (typically featuring two class II PhaCs) being notably conserved (Ciesielski et al., 2014; Mozejko-Ciesielska et al., 2019), our results showed that not every strain is capable of PHA production, with some lacking PhaC genes. Nevertheless, the genus garners significant attention as it is one of the most important bacteria for PHA manufacturing at an industrial scale (Poblete-Castro et al., 2020; Weimer et al., 2020). In this line, 86% of the *Pseudomonas* strains analyzed encoded two class II PhaCs, suggesting they would have been present in the *Pseudomonas* last common ancestor and then inherited vertically and subjected to speciation-driven divergence. Notably, besides the PhaC genes belonging to the core genome of most *Pseudomonas* species, we found several cases of strains carrying genes encoding additional class II PhaCs. Moreover, we identified 42 strains from 25 species encoding class I PhaCs, a finding until now considered rare in this genus (Ayub et al., 2007; Mozejko-Ciesielska et al., 2019).

Several strains showing additional PhaC genes corresponded to Antarctic isolates, highlighting the particular capacities of members of this genus thriving in extreme environments (Ayub et al., 2007; Tan et al., 2020). As a notable example, the Antarctic strain *P. frigusceleri* MPC6 encodes five PhaCs, three class II, one class I, and one markedly distant from all known PhaC classes, named PhaC3 (Orellana-Saez et al., 2019). Although conserved domain analysis suggests that PhaC3 may be closer to class I PhaCs, phylogenetic analysis revealed that it substantially diverges from all the class I PhaCs in our database, suggesting it would represent a novel class. Notably, MPC6 was the only strain from our genome dataset bearing this PhaC class. Nevertheless, an extended BLASTp search against the NCBI database uncovered four *Pseudomonas* spp. strains encoding proteins sharing high identity (>95%) with PhaC3. The respective genes were encoded in putative genomic islands integrated into tRNA-Sec genes. These results indicate that, while rare, PhaC3 is not unique to MPC6, with presence in other *Pseudomonas* species.

Our discovery of PhaC3 aligns with a report of four Antarctic bacterial strains with uncommon traits: the presence of a class I PhaC within the *Pseudomonas* genus and a novel unclassified PhaC within *Janthinobacterium* that diverges from the known classes (Tan et al., 2020). These insights suggest that Antarctica may be a source of previously unidentified PhaCs and that Class I PhaCs within *Pseudomonas* would be more frequent than previously thought. In this regard, PHA-producing microorganisms usually benefit from having this polymer in the cell under fermenting process and environmental stress conditions, typical of Antarctic environments, suggesting an adaptive role for PHA synthesis in *Pseudomonas* strains (Borrero-de Acuña et al., 2021; Obruca et al., 2018; Spiers et al., 2000). *P. frigusceleri* MPC6 demonstrates adaptability with its five PhaCs, capable of utilizing an ample range of substrates to synthesize both block or homopolymers of scl-and-mcl-PHA depending on the carbon source used, with comparable yields across a wide temperature range (Orellana-Saez et al., 2019; Pacheco et al., 2019). Additionally, gene duplication is a hallmark of positive selection in evolution, often conferring significant advantages (Ohno, 1970). Thus, an expanded PhaC set would increase fitness, especially in environments with limited nutrient availability where metabolic versatility is required.

The current consensus for the classification of PhaC enzymes (type I to IV) is based on three criteria: i) substrate specificity *in vivo*, ii) the constitutive subunits (PhaC, PhaE, or PhaR), and iii) amino acid sequence (Rehm, 2003). Nevertheless, the discovery of atypical PhaCs within *Pseudomonas* species and the increasing known diversity of these enzymes challenges the adequacy of current classification systems. As an additional challenge, high amino acid sequence diversity is found within current PhaC classes (Tan et al., 2020). Moreover, another caveat is that, although it is accepted that class I, III, and IV PhaCs can only incorporate scl-PHA monomers and class II only mcl-PHA, a study pointed out that there are exceptions where PhaCs closely resembling class I enzymes that can incorporate mcl-PHA (Neoh et al., 2022). In the same study, PhaC1 and PhaC2 of *Pseudomonas sp.* 61-3 are highlighted due to their high similarity to class II PhaCs and their ability to incorporate scl-PHA monomers forming scl-*co*-mcl-PHA copolymers. Hence, the bioinformatics dimension must still be better incorporated into the PhaCs classifications.

Regardless of the great importance of PHA synthesis for biotechnological purposes and the key PhaC role in this process, only two PhaCs have an experimentally resolved structure of their catalytic domains, one from *C. necator* and one from *Chromobacterium sp*. (Neoh et al., 2022). This limitation hampers structural comparisons among different PhaCs, especially in other bacterial lineages, including *Pseudomonas*. To progress in this direction, we leveraged the recent developments in artificial intelligence-driven protein structure modeling to generate 3D structure models for representatives of all the PhaC classes in our *Pseudomonas* set. Despite the high PhaC sequence diversity inside and between different classes, all showed remarkable structure similarity. Also, all had the conserved N-terminal domain phaC_N (pfam07167). However, the exact function of this domain is still unclear, as is the impact of variations in length between different PhaCs on its function (Neoh et al., 2022). Even among different classes (I, II, and X), which share very low amino acid sequence identity, the global topology is conserved even beyond *Pseudomonas*. For instance, class X PhaC from *P. frigusceleri* showed a high structural similarity with class I enzyme from *C. necator*.

Phylogenomic analyses revealed deep-branched lineages among the *Pseudomonas* strains (except for *P. aeruginosa*), highlighting the marked genetic heterogeneity inside this bacterial group and several incongruences in species classification that should be reviewed. A clear example corresponds to *P. fluorescens*, the most diverse group within the genus *Pseudomonas*, which should be considered a species complex (Garrido-Sanz et al., 2017). In agreement, we observed *P. fluorescens* isolates distributed in different tree positions. This situation was also observed in other species, including *P. putida*.

The sparse distribution of PhaC genes other than those belonging to the canonical cluster, with variations even among closely related strains, strongly suggested their acquisition by horizontal transfer. Indeed, genomic context analysis of sporadic PhaC genes revealed that most are associated with mobile genetic elements, including plasmids, phages, and genomic islands. A few previous reports indicate that PhaC genes may be transferred via horizontal transference. One such study revealed that the Antarctic strain *Pseudomonas* sp. 14-3 harbors a 32.3-kbp genomic island named pha-GI, which carries a *phaC* gene and different genes and elements associated with mobility, including an integrase and insertion sequences (Ayub et al., 2007). Moreover, a follow-up study with this strain explored the effect of PHA accumulation on the adaptability to cold conditions using a PHA synthase-minus mutant (Δ*phaC*). The study revealed that PHAs are essential for maintaining the redox state in this Antarctic bacterium during adaptation to low temperatures (Ayub et al., 2009). Considering that many reported Antarctic *Pseudomonas* have additional PhaC genes, it is plausible that the extreme environment could establish a selective pressure toward promoting PHA synthesis and make it more versatile. Nevertheless, some Antarctic *Pseudomonas* lacked PhaC genes, while some isolates from other environments (including host-associated *P. aeruginosa*) showed increased PhaC gene content. Additional studies are required to understand better PHA’s diverse roles in different bacteria and niches.

Other evidence of PhaC genes likely acquired through horizontal transfer and their impact on PHA production includes findings in *P. extremaustralis*, where a *phaC* gene presumably acquired horizontally showed higher expression compared to the native mcl-PHA synthases, conferring the ability to produce high amounts of PHB (Brito et al., 2024; Catone et al., 2014). Moreover, the *phaC* genomic region showed high similarity with a genomic island described in *P. stutzeri* A1501. Additionally, *P. umsongensis* GO16 was shown to accumulate scl-PHAs, which is considered not a common characteristic of *Pseudomonas* bacteria. However, this ability was demonstrated for various *Pseudomonas* species, including *P. oleovorans*, *P. pseudoalcaligenes*, *Pseudomonas* sp., as well as recently described *P. extremaustralis* and *Pseudomonas frigusceleri* MPC6 (Narancic et al., 2021; Orellana-Saez et al., 2019). This ability was linked to an expanded set of PhaC genes likely acquired horizontally.

Beyond *Pseudomonas*, a previous study investigating the diversity and phylogenomics of the PHA biosynthesis based on the analysis of 253 genomes, found that discrepancies in the phylogenetic trees for *phaA*, *phaB,* and *phaC* genes suggest that HGT may be a major contributor to its evolution (Kalia et al., 2007). This study found *phaC* genes in 40 genera from diverse taxonomic groups, such as Actinobacteriota, Cyanobacteriota, Bacillota, Pseudomonadota, and Eukaryotes. In this same direction, horizontal gene transfer of genes to produce PHAs and ectoine has been observed in *Halomonas* sp. TD01, with the presence of potentially novel classes of these enzymes, highlighting the caveats of the current PhaC classification and the potentially vast PhaC diversity yet to be discovered (Cai et al., 2011).

## CONCLUSIONS

Overall, we demonstrated a significant diversity of genes encoding PhaC enzymes in *Pseudomonas*, encompassing a variety of class II and I subgroups and even enzymes belonging to a potentially novel class. Despite notable differences in amino acid sequence, these proteins exhibit remarkable structural conservation, sharing a highly similar topology, secondary structure elements, and conserved domains. However, the encoding genes are highly variable and, in many cases, located in different types of mobile genetic elements, including plasmids, phages, and genomic islands. Therefore, PhaC genes would be frequent substrates for horizontal gene transfer and, thus, likely under positive selection. These results highlight the remarkable metabolic plasticity of some *Pseudomonas* strains, particularly those adapted to thrive in extreme environments like Antarctica. Future studies are required to unveil the functional implications of the PhaC diversity on PHA synthesis and its biotechnological applications.

## DATA AVAILABILITY

The data underlying this article were accessed from the NCBI Genome Database (https://www.ncbi.nlm.nih.gov/home/genomes). The accession numbers of the genomes are provided in Table S1. The derived data generated in this research will be shared on reasonable request to the corresponding author.

## AUTHORS CONTRIBUTIONS

Cox-Fermandois A. (Conceptualization, Data curation, Formal analysis, Investigation, Methodology, Visualization, Writing - original draft, Writing - review & editing), Berríos-Pastén C. (Data curation, Formal analysis, Investigation, Methodology, Validation, Visualization), Serrano C (Formal analysis, Methodology, Validation, Visualization), Arros P (Data curation, Formal analysis, Investigation, Validation, Visualization), Poblete-Castro I (Conceptualization, Investigation, Supervision, Validation, Visualization, Writing - original draft, Writing - review & editing), Marcoleta AE (Conceptualization, Data curation, Formal analysis, Funding acquisition, Investigation, Methodology, Project administration, Resources, Supervision, Validation, Visualization, Writing - original draft, Writing - review & editing).

