## Supplementary material for "*Pseudomonas* species from Antarctica and other environments host diverse genes encoding structurally conserved polyhydroxyalkanoate synthases linked with different mobile genetic elements"

### Large-scale analysis of polyhydroxyalkanoate synthases in Pseudomonas: highly diverse enzymes with potential for a novel class and dissemination by horizontal gene transfer

Amelia Cox-Fermandois[1], Camilo Berríos-Pastén[1], Carlos Serrano[1], Patricio Arros[1], Ignacio Poblete-Castro[2], and Andrés Marcoleta[1,*]

[1]Grupo de Microbiología Integrativa, Facultad de Ciencias, Universidad de Chile. Las Palmeras 3425 Ñuñoa, Santiago, Chile.

[2]Biosystems Engineering Laboratory, Department of Chemical and Bioprocess Engineering, Faculty of Engineering, Universidad de Santiago de Chile (USACH), 9170022, Santiago, Chile

#### SUPPLEMENTARY MATERIAL

**Figure S1.** Heatmap showing the pairwise identity between class-I and class-X enzymes from the different sub-classes found. The presence of conserved domains is indicated for each protein.

**Figure S2.** Class I.4 PhaC genes in *P. oleovorans* Ppseudo_Pac and CECT5344 are encoded in a putative genomic island integrated into a Thr tRNA gene.

**Figure S3.** Representatives class I PhaC genes from different subclasses showing a set of genes typically found linked to and sharing their genomic context with PhaC genes.

**Table S1.** *Pseudomonas* genome set analyzed in this study (provided as a separate spreadsheet).

**Table S2.** Protein sequences used as references to search for PhaC enzymes of different classes among *Pseudomonas* genomes.

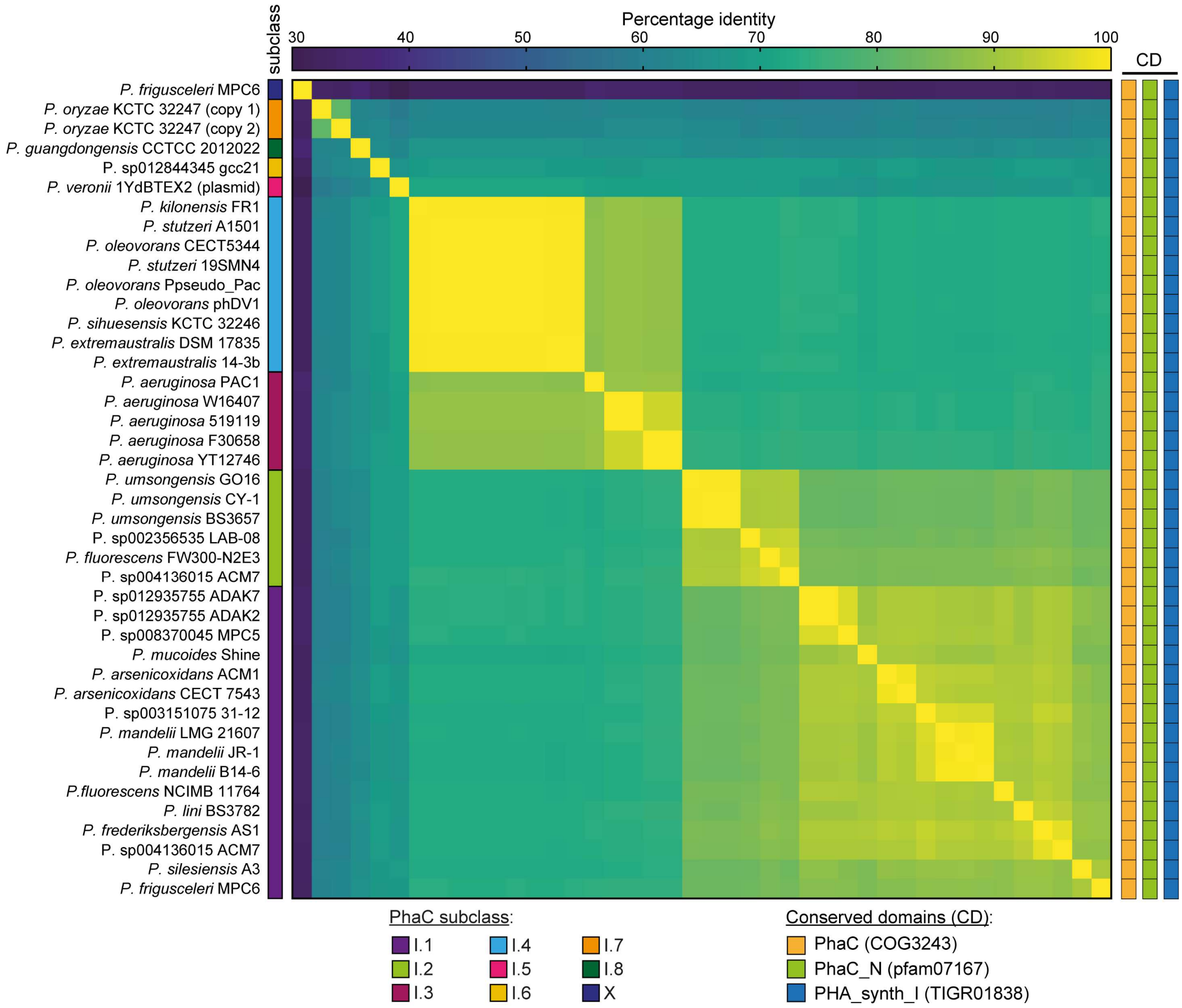


**Figure S1.** Heatmap showing the pairwise identity between class-I and class-X enzymes from the different sub-classes found. The presence of conserved domains is indicated for each protein.

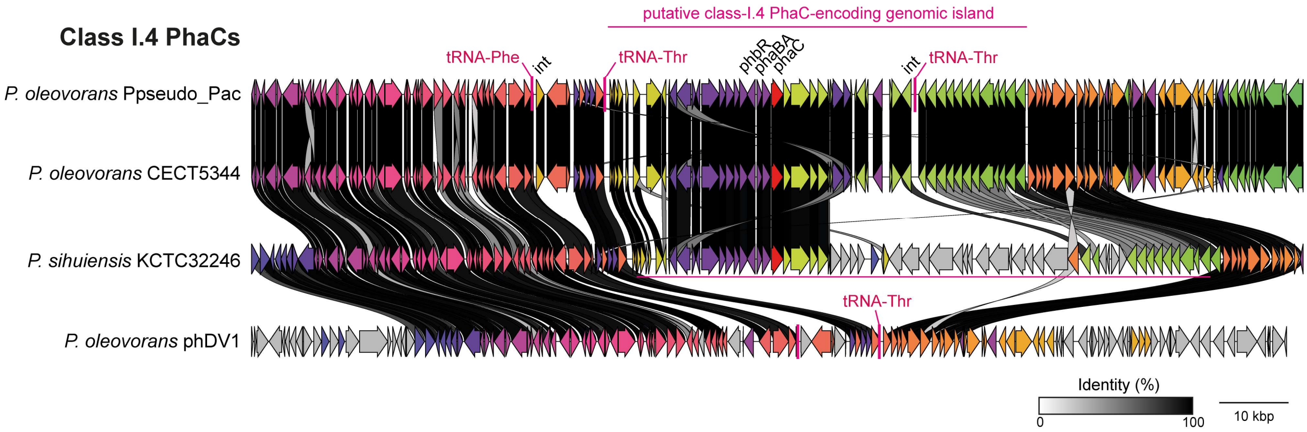


**Figure S2.** Class I.4 PhaC genes in *P. oleovorans* Ppseudo_Pac and CECT5344 are encoded in a putative genomic island integrated into a Thr tRNA gene. The strains *P. sihuiensis* KCTC32246 and *P. oleovorans* phDV1 lacking class I.4 PhaC genes were included for comparison.

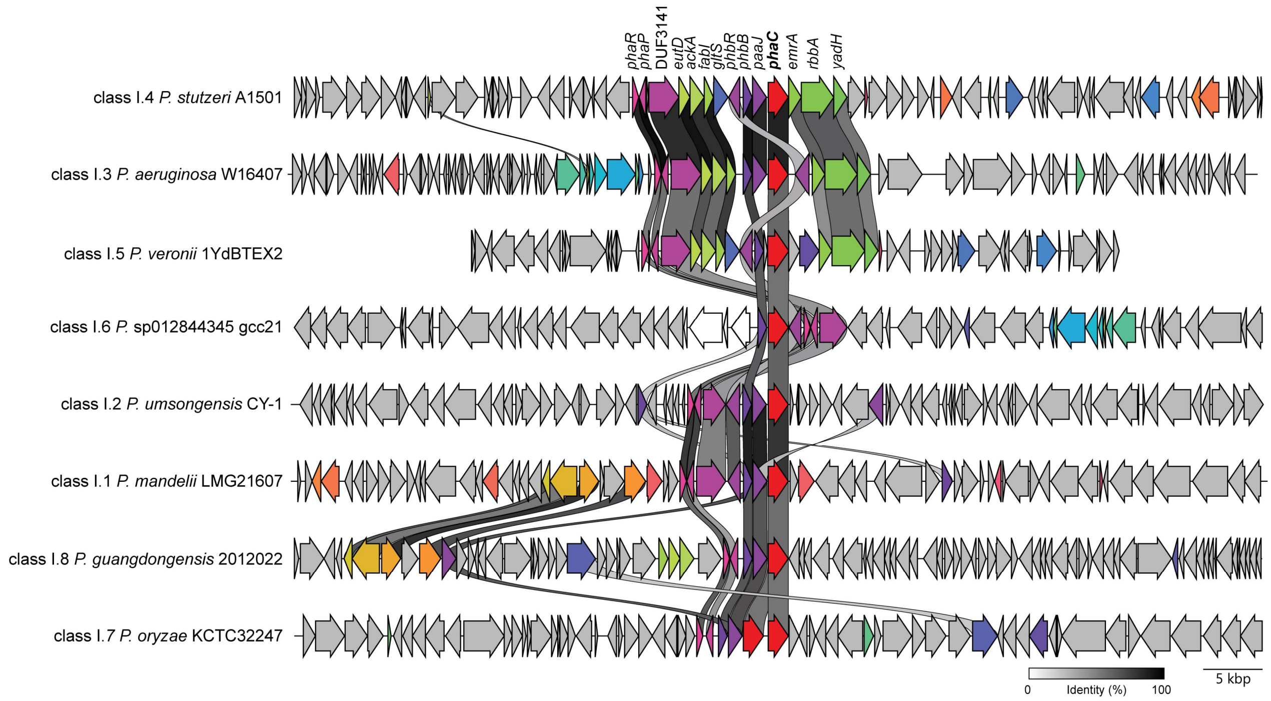


**Figure S3.** Representatives class I PhaC genes from different subclasses showing a set of genes typically found linked to and sharing their genomic context with PhaC genes.

**Table S2.** Protein sequences used as references to search for PhaC enzymes of different classes among *Pseudomonas* genomes.

| PhaC class | NCBI accession | Source organism |
|---|---|---|
| Class I | WP_011615085.1 | *Cupriavidus necator* |
| Class II | ACK57560.1 (PhaC1) | *Pseudomonas fluorescens* |
| | ACK57562.1 (PhaC2) | *Pseudomonas fluorescens* |
| | AAM63407.1 (PhaC1) | *Pseudomonas putida* |
| | AAM63409.1 (PhaC2) | *Pseudomonas putida* |
| | AAG08441.1 (PhaC1) | *Pseudomonas aeruginosa* |
| | AAG08443.1 (PhaC2) | *Pseudomonas aeruginosa* |
| Class III | ACB10370.1 (PhaC) | *Haloferax mediterranei* |
| | ACB10369.1 (PhaE) | *Haloferax mediterranei* |
| Class IV | AAD05260.1 (PhaC) | *Priestia megaterium* |
| | AAD05258.2 (PhaR) | *Priestia megaterium* |
